# Population genomics of *Xenopus laevis* in southern Africa

**DOI:** 10.1101/2022.07.21.501003

**Authors:** Tharindu Premachandra, Caroline M. S. Cauret, Werner Conradie, John Measey, Ben J. Evans

## Abstract

Allotetraploid genomes have two distinct genomic compartments called subgenomes that are derived from separate diploid ancestral species. Many genomic characteristics such as gene function, expression, recombination, and transposable element mobility may differ significantly between subgenomes. To explore the possibility that subgenome population structure and gene flow may differ as well, we examined genetic variation in an allotetraploid frog – the African clawed frog (*Xenopus laevis*) – over the dynamic and varied habitat of its native range in southern Africa. Using reduced representation genome sequences from 91 samples from 12 localities, we found no strong evidence that population structure and gene flow differed substantially by subgenome. We then compared patterns of population structure in the nuclear genome to the mitochondrial genome using Sanger sequences from 455 samples from 183 localities. Our results provide further resolution to the geographic distribution of mitochondrial and nuclear diversity in this species and illustrate that population structure in both genomes corresponds roughly with variation in seasonal rainfall and with the topography of the southern Africa.

## Introduction

### Polyploidization

Whole genome duplication (WGD or polyploidization) can occur spontaneously within a species (autopolyploidization) or in association with hybridization among different species (allopolyploidization). Polyploidization preceded extraordinary diversifications such as eudicot plants, jawed vertebrates, and teleost fish, and more recently in many important species (salmonids, wheat, corn, cotton, rice, yeast, paramecium, tetrahymena, and more), raising the questions of how WGD influences adaptation, diversification, and genome evolution (Otto and Whitton 2000; Van de Peer et al. 2009; Bomblies 2020; Lovell et al. 2021).

Polyploid genomes are distinguished from diploid genomes in gene copy number, but also in several other ways. For example, tetrasomic inheritance – where four alleles are present at individual loci – may be associated with chromosomal mis-segregation, favoring genetic variation that leads to diploid-like (disomic) meiosis (Soares et al. 2021). Disomic inheritance can be achieved when recombination becomes rare or ceases between each compartment or “subgenome” of a polyploid genome that is derived from one (autopolyploids) or two (allopolyploids) ancestral species, thereby allowing the subgenomes to evolve independently with little or no inter-subgenome recombination. Subgenomes thus may evolve distinctive features such as differences in gene expression, gene silencing, alternative splicing, rates of recombination and rearrangement, rates of protein evolution, and complements and mobilities of transposable elements and other repetitive elements (Liu et al. 2014; Session et al. 2016; Elurbe et al. 2017; Mei et al. 2017; Cheng et al. 2018; Furman et al. 2018; Edger et al. 2019; Lee and Adams 2020; Yu et al. 2020; Zhao et al. 2020; Schiavinato et al. 2021). In allopolyploids, duplicated genes may each have disomic inheritance upon formation, owing to divergence between pairs of homeologous chromosomes (reviewed in Wolfe 2001).

Further complexity arises when subgenomes establish disomic inheritance at different times: prior to the origin of disomic inheritance, genetic drift or natural selection can contribute to the elimination of variation from one ancestral species in portions of the genome with polysomic inheritance (Wolfe 2001). However, genetic exchanges between subgenomes such as translocation, exchange of transposable elements, and recombination events such as gene conversion may occur. For example, in the allopolyploid frog *Xenopus laevis*, a sex determining gene called *dm-W* resides in one subgenome (the “L” subgenome), but is derived from a partial duplication of a gene (*dmrt1S*) that resides in the other subgenome (the “S” subgenome; Bewick et al. 2011).

### Subgenome asymmetry in gene flow and population structure

Within each subgenome, gene flow among populations may differ. Consider, for example, a hypothetical disomic allotetraploid individual that is a progeny of a cross between two parental allotetraploid populations (Fig. 1). This individual would be heterozygous in both subgenomes for population-specific variation from each parental population. A gamete of this individual thus would carry a mosaic of alleles from each parental population, with 50% probability of carrying a population-specific allele from either one. If backcrossed to an individual from one parental population, the resulting backcrossed individuals would have a mosaic of homozygous and heterozygous genotypes from both parental populations. This genetic variation among siblings is subject to natural selection and genetic drift, which could result in asymmetry in the extent of introgression in each subgenome if backcrossed survivors join a parental population (an example of more gene flow in subgenome 2 than subgenome 1 is depicted in Fig. 1).

**Fig. 1.**
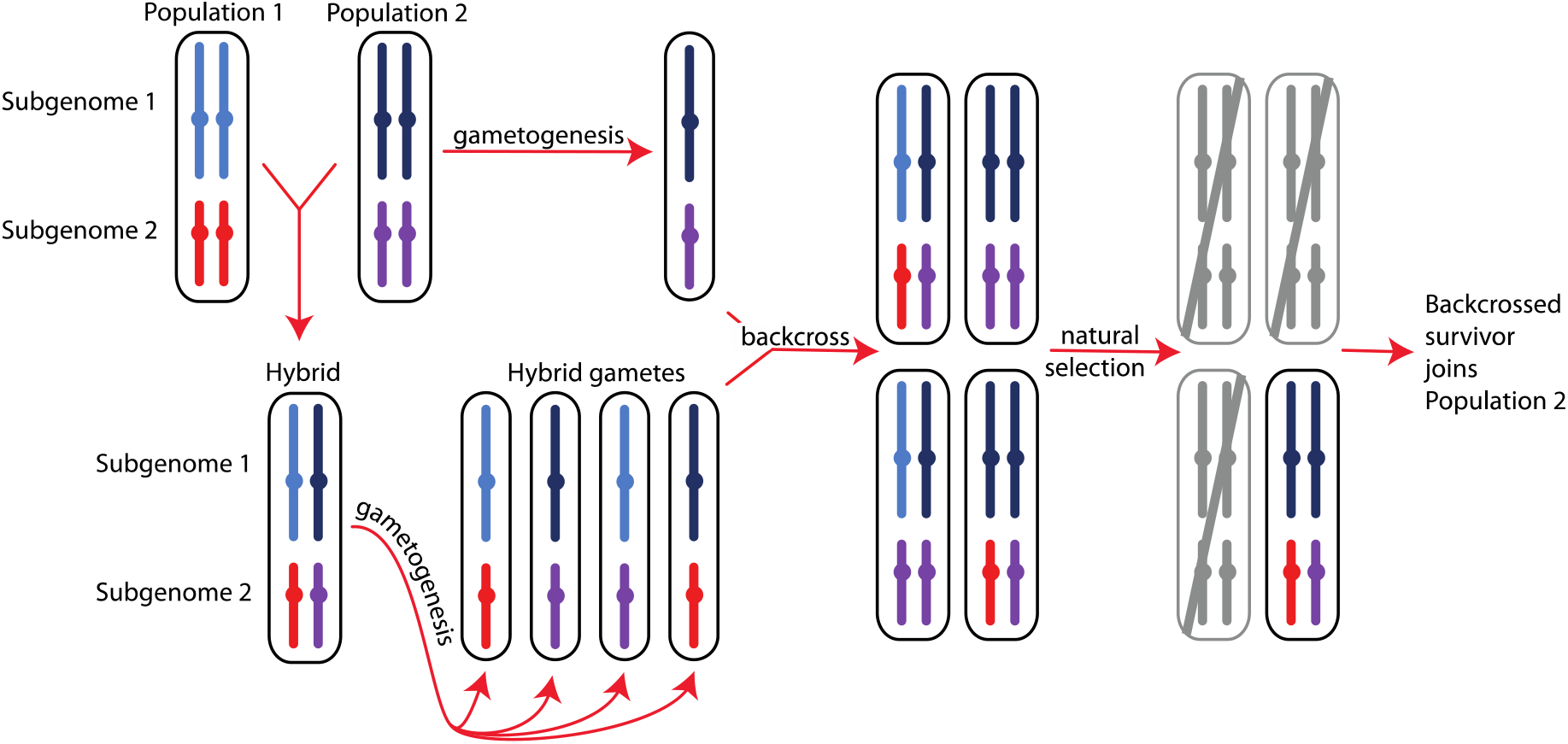
Differential introgression in subgenomes can be achieved randomly or in association with natural selection against some admixed genotypes as shown above. The parental populations (Population 1 and 2) are assumed to have two subgenomes with disomic inheritance (indicated with light blue and dark blue or red and purple, respectively, with color differences within each subgenome representing population subdivision). Gametogenesis in a hybrid individual produces gametes with a mosaic of population variation in each subgenome, depicted here as blocks that include entire chromosomes (in reality, recombination would likely generate smaller blocks of population-specific variation within each chromosome). If some backcrossed individuals have low fitness (gray) and high fitness individuals successfully breed in one parental population, different proportions of each subgenome may introgress, here shown as an extreme example of introgression only in subgenome 2.

Because cytoplasmic genomes are generally not duplicated during polyploidization events, asymmetry in subgenome evolution has the potential to be influenced by the species of each hybrid cross which was maternal (Sharbrough et al. 2021). Thus, not only might patterns of subgenome evolution differ, but also population genetic and phylogenetic patterns in the mitochondrial genome may differ from one or both subgenomes.

### Polyploid ratchet

Gene loss, gene conversion, and other forms of recombination may erode or homogenize variation within each subgenome. These genomic changes can have fitness consequences, such as centromere homogenization and distorted segregation, that create a downward fitness spiral called the “polyploid ratchet” where genomic instability begets further instability (Gaeta and Pires 2010). The polyploid ratchet is supported by comparisons between natural and synthetic polyploids: rearrangements in synthetic allopolyploids are frequently more common than in (older) natural allopolyploids, suggesting the action of purifying selection and other processes that increase genomic stability over time (Gaeta et al. 2007). For example, subgenome-specific gene flow (between populations or species) has the potential to counteract the polyploid ratchet by restoring genetic variation that was lost or damaged during allopolyploid evolution. In allotetraploid barbed goatgrass (*Aegilops triuncialis*), for example, gene flow with domestic wheat is higher in one subgenome compared to the other, which is possibly linked to variation between subgenomes in genomic stability and gene content (Parisod et al. 2013). Differential gene flow in each subgenome also has potential phenotypic consequences. In various plants from the agriculturally important genus *Brassica*, several allotetraploids are derived from different combinations of three diploid progenitors; subgenome-specific introgression between these allotetraploids and other synthetic polyploids corresponds with variation in phenotypes (Wei et al. 2016).

### Allopolyploid evolution in African clawed frogs

African clawed frogs (genus *Xenopus*) underwent at least two independent allotetraploidization, three independent allooctoploidization, and three independent allododecaploidization events, which gave rise to a striking diversity of polyploid species: 13 allotetraploids, seven allooctoploids, at least three allododecaploids (Evans 2008; Evans et al. 2015). A probable mechanism for allopolyploidization involves successive generations of backcrossing and production of unreduced gametes by hybrid females, and has been recapitulated in crosses of captive animals (Kobel and Du Pasquier 1986). Within the species *Xenopus laevis*, subgenomes are karyotypically distinguished by distinctive sizes of homeologous chromosomes, such that the “L” and “S” subgenomes have “long” and “short” versions of homeologous chromosomes (Matsuda et al. 2015; Session et al. 2016). A striking finding that emerged from analysis of a high quality, chromosome-scale genome sequence of *X. laevis* (Session et al. 2016; Elurbe et al. 2017), and other allotetraploid *Xenopus* (Furman et al. 2018) is that each subgenome also has distinctive molecular genetic characteristics, with the L subgenome encoding transcripts that have a lower rate of gene silencing, higher expression level, stronger purifying selection, and longer coding regions compared to homeologous transcripts from the S subgenome. Additionally, the S subgenome experienced more rearrangements than the L subgenome (Session et al. 2016).

### Xenopus laevis as a model organism for biology

*Xenopus laevis* has an unusual relationship with humans, having first been deployed as a pregnancy test (Weisman and Coates 1941), which led to its widespread export from a portion of its native range in the southwestern South Africa, to Europe, North America, and Asia (Gurdon and Hopwood 2000; Weldon et al. 2007; van Sittert and Measey 2016). By the mid-20^th^ century, this species had been adopted as a model organism for developmental biology (Cannatella and de Sá 1993; Gurdon 1996; Harland and Grainger 2011) owing in large part to its relative ease of maintenance in captivity and other useful attributes such as external fertilization and development. *Xenopus laevis* is also prevalent in the pet trade (Measey 2017) and invasive populations are now established in the United States, Mexico, Chile, France, Italy, Portugal, Japan, and China (Rebelo et al. 2010; Measey et al. 2012; Lillo et al. 2013; Peralta-Garcia et al. 2014; Ihlow et al. 2016; Wang et al. 2019; Ginal et al. 2021; Vimercati et al. 2021). Historical records demonstrate that exported individuals were sourced from multiple localities in southern Africa (van Sittert & Measey 2016), and some invasive populations are admixed (De Busschere et al. 2016; Measey et al. 2017).

### Taxonomy and molecular phylogeography of X. laevis

*Xenopus laevis* was described over 200 years ago (Daudin 1802) and has an extensive taxonomic history (Frost 2021). Previously, *X. laevis* included several subspecies (*X. l. laevis, X. l. victorianus, X. l. petersii, X. l. poweri, X. l. sudanensis, X. l. bunyoniensis*) with a massive distribution over much of sub-Saharan Africa (Kobel et al. 1996; Tinsley et al. 1996). However, based on an analysis of molecular variation in mitochondrial and nuclear markers, it was proposed that *X. laevis sensu lato* (Kobel et al. 1996; Tinsley et al. 1996) comprised multiple species, including *X. laevis*, (southern Africa), *X. victorianus* (East Africa), *X. petersii* (west central Africa), and *X. poweri* (northwest central Africa; Furman et al. 2015). Within *X. laevis sensu stricto* (Furman et al. 2015), hereafter *X. laevis*, mitochondrial DNA is paraphyletic, and considerable population structure exists, especially in the southwest Cape Region on either side of the Cape Fold Mountains (Grohovaz et al. 1996; Evans et al. 1997; Measey and Channing 2003; Evans et al. 2004; Du Preez et al. 2009; Furman et al. 2015).

The distribution of *X. laevis* spans multiple biomes including (from southwest to northeast): fynbos, succulent Karoo, Nama-Karoo, forest, thicket, savanna, and grassland (Mucina and Rutherford 2006), and patterns of population structure roughly corresponds with geographic variation in annual rainfall patterns with winter rainfall in the southwest and summer rainfall in the northeast (Grohovaz et al. 1996; Evans et al. 1997; Measey and Channing 2003; Evans et al. 2004; Du Preez et al. 2009; Furman et al. 2015). Phenotypic variation exists within *X. laevis*, such as size differences (larger in the SW) and variation in gonadal morphology, with a higher frequency of testicular ovarian follicles detected in animals originating from northeast compared to southwest South Africa (Du Preez et al. 2009). Reciprocal translocation experiments suggest adaptation to different rainfall and altitudinal regimes, as well as variation in the extent of plasticity of tadpoles to adjust to these regimes (Wagener et al. 2021; Kruger et al. 2022). Overall, the substantial and complex population structure of *X. laevis* coupled with its wide use as an experimental organisms and new role as an invasive species, underscores the importance of understanding genetic variation in natural populations and how this relates to diversity in captive and invasive populations.

*Goals. Xenopus laevis* is a particularly promising species for exploration of distinctive patterns of subgenome introgression because the subgenomes have different degrees of genomic instability, because this species resides in a diverse landscape, and because a high quality genome assembly is available that allows one to identify subgenome-specific genetic variation and quantify subgenome-specific gene flow. We had an *a priori* expectation that, across the native habitats of *X. laevis*, the L subgenome would have more geographic structure (less gene flow) than the S subgenome. This expectation derives from (i) previous observations of stronger purifying selection on genes (Furman et al. 2018) and a lower rate of pseudogenization (Session et al. 2016) in the L subgenome, presumably including some genes that are involved with adaptation to local ecological conditions, and (ii) previous observations of higher genomic instability in the S subgenome (Session et al. 2016), which could be associated with the polyploid ratchet and potentially mitigated by gene flow. To test this expectation, we set out to further characterize genetic variation in each *X. laevis* subgenome in its natural range with a focus on southwestern South Africa where most of the genetic variation occurs, including samples within and on either side of an intraspecific contact zone (Du Preez et al. 2009; Furman et al. 2015) that corresponds to changes in both biome and rainfall (Chase and Meadows 2007). We also compared patterns of genetic variation in each subgenome based on reduced representation genome sequences (RRGS) data to that in the mitochondrial genome based on Sanger sequences from an even more widespread sample of wild caught animals.

## Methods

### Samples and genetic data

A survey of genetic variation in the nuclear genome was carried out using RRGS from a panel of 90 wild caught *X. laevis* individuals from 12 localities (Table S1; Fig. 1), one *X. victorianus* from the Democratic Republic of Congo, three *X. poweri* samples (one each from Nigeria, Cameron, and Botswana), and one *X. gilli* sample from South Africa. Genetic variation in the mitochondrial genome was characterized using Sanger sequencing of a ∼814 base pairs portion of the mitochondrial 16S rRNA gene in a total of 455 samples from 183 localities (Fig. 1), including 374 *X. laevis* samples from 183 localities (of which 105 were previously published; Furman et al. 2015), one *X. gilli* sequence that we defined as an outgroup based on previous analyses (Evans et al. 2004; Evans et al. 2019), and 41, 23, and 16 samples of *X. victorianus, X. poweri*, and *X. petersii* samples (of which 41, 23, and 7 respectively were previously published; Furman et al. 2015). We used primers 16Sc-L and 16Sd-H (Evans et al. 2003) for amplification and sequencing.

For the RRGS data, normalization and library preparation was performed at the Genomic Centre at the University of Laval using a double digest restriction enzyme associated DNA protocol with SbfI and MspI restriction enzymes. The libraries were run on one third of lane of a NovaSeq S4 machine with paired end 150 base pair reads. Reads were demultiplexed using Sabre (Joshi 2011), and trimmed using Cutadapt (Martin 2011) and Trimmomatic version 0.39 (Bolger et al. 2014).

RRGS reads were aligned to the *X. laevis* genome assembly version 9.2 (Session et al. 2016) using the mem function of Bwa version 0.7.17 (Li and Durbin 2009) and Samtools version 1.10 (Li et al. 2009). Readgroups were added with Picard version 2.23.3 (Development_team) and alignment and genotyping with the Genome Analysis Toolkit (GATK) version 4.1.0.0 (McKenna et al. 2010) using the HaplotypeCaller, GenomicsDBImport, and GenotypeGVCFs functions. Data were then hard filtered using the VariantFiltration and SelectVariants functions of GATK with filtering criteria recommended by best practices guide of the Broad Institute. Positions with the following attributes were removed: QD > 2.0, QUAL < 20, SOR > 3.0, FS > 60.0, MQ < 40.0, MQRankSum < -12.5, and ReadPosRankSum < -8.0, where these acronyms refer to variant confidence/quality by depth (QD), genotype quality (QUAL), Symmetric Odds Ratio of 2×2 contingency table to detect strand bias (SOR), Fisher exact test for strand bias (FS), map quality (MQ), a test for read map quality (MQRankSum), and a test for read position bias (ReadPosRankSum), respectively.

### Analyses of RRGS data

Geographical structure of genetic variation in each subgenome of *X. laevis* was visualized using principal components analysis (PCA) using the SNPRelate (Zheng et al. 2012) package and Admixture (Alexander et al. 2009). These analyses were performed separately for variable positions in each subgenome using the RRGS data mapped to the whole genome after excluding unplaced scaffolds and excluding samples that were not *X. laevis*. For the principal components analysis, SNPs were pruned with a linkage disequilibrium threshold of 0.2 and requiring at 50% of the samples to have called genotypes. After pruning, a total of 59,998 and 47,203 positions were considered for the L and S subgenomes, respectively. Prior to Admixture analysis the data were thinned by requiring at least 50% of the samples have called genotypes.

We quantified genetic exchange across an area where winter rainfall changes to summer rainfall, and biomes change from fynbos and succulent karoo to Nama-karoo (Chase and Meadows 2007) using Patterson’s *D* statistic (Patterson et al. 2012). Patterson’s *D* is calculated between three ingroup populations (P_1_, P_2_, P_3_) and an outgroup (O) whose evolutionary relationships follow this topology: (((P_1_, P_2_), P_3_), O). Positive values indicate an excess of shared sites between P_2_ and P_3_ (ABBA sites) whereas negative values indicate an excess of shared sites between P_1_ and P_3_ (BABA sites). We calculated Patterson’s *D* using scripts in the genomics_general package (Martin 2021) with four non-independent parameterizations (Comparisons 1–4) that were based on patterns of genome wise similarity in the RRGS data discussed below. In addition, we performed a focused analysis on admixed individuals from Laingsburg relative to adjacent populations (Beaufort West, De Doorns, and Victoria West + Kimberley). Confidence intervals for Patterson’s *D* were obtained using the block jackknife (Green et al. 2010). Patterson’s *D* was calculated for each subgenome using the full dataset in using 5 Mb genomic windows and using a subset of each subgenome that included only genic regions only (i.e., introns and exons) in 10 Mb genomic windows.

A permutation test was used to evaluate the one-sided expectation that Patterson’s *D* statistic should be more extreme for the S than the L subgenome. As discussed above, this expectation derives from the possibility of more locally advantageous variation in the L subgenome and the possibility of restorative gene flow in the S subgenome that could counteract the polyploid ratchet.

The test statistic of the permutation tests was the difference in the average Patterson’s *D* statistic for each genome, which was calculated with weighting by the sum of the number of ABBA and BABA sites in each genomic window. This test statistic was compared to the distribution of 1000 statistics that were obtained by permutating the L and S subgenome assignments of each window and then re-calculating the difference between the weighted averages.

Because the permutation tests generally found no significant difference between the L and S subgenome, we evaluated the power of our data to detect a difference. For each comparison, the standard error of D of each subgenome was calculated following Busing et al. (1999). Quantile-quantile plots suggested that the distributions of Patterson’s *D* within genomic blocks were approximately normally distributed within each subgenome (data not shown). The variance of Patterson’s *D* was therefore estimated as the product of the squared standard error and the number of genomic blocks in each subgenome. The standard error of the test statistic is thus the square root of the sum of the within subgenome variances divided by the number of windows in each subgenome (Swinscow and Campbell 2002). Using this value we calculated an estimate of the test statistic where one would expect an 80% probability of detecting a significant difference, which is equal to 2.8 times the pooled standard error of the test statistic (Gelman and Hill 2006). For each comparison, we also calculated the Cohen’s d effect size (Cohen 2013), which is the difference between the mean divided by the pooled standard deviations, which we estimated as the square root of the average variance in the L and S subgenomes. The effect size measures the strength of the relationship between two variables (here Patterson’s *D* and subgenome); values below 0.2 indicate a weak relationship (Cohen 2013).

### Phylogenetic analysis of mitochondrial and nuclear DNA

*Phenograms*. Because the population genetic relationships among the *X. laevis* autosomal data violate assumptions of a strictly bifurcating relationship among individuals, we summarized genetic distances between individuals using a neighbor-joining tree which was calculated using Jukes-Cantor distances (Jukes and Cantor 1969) in PAUP* (Swofford 2002). An input (nexus) file was generated from the genotypes using the vcf2phylip script (Ortiz 2019) and confidence of nodes was evaluated using 1,000 bootstrap replicates. We performed this analysis using variable positions only and requiring at least 80 out of the 95 samples to have called genotypes. This resulted in datasets with 1,018,084 and 801,859 variable positions for the L and S subgenomes, respectively.

We estimated evolutionary relationships among the partial mitochondrial sequences using the maximum likelihood criterion and the software IQ-TREE version 1.6.12 (Nguyen et al. 2015). The GTR+F+I+G4 model was used based on the Bayesian Information Criterion as implemented by IQ-TREE; confidence of nodes was evaluated using the ultrafast bootstrap approach (Minh et al. 2013). Additionally for the partial mitochondrial sequences, we constructed a parsimony network using TCS (Clement et al. 2000) and visualized with PopArt (Leigh and Bryant 2015).

### Data availability

The RRGS data from this study are listed in Table S1 and have been deposited in the Short Read Archive of NCBI (BioProject TBA). All new Sanger sequences are listed in Table S2 and have been submitted to GenBank (accession nos TBA-TBA).

## Results

### PCA, Admixture

Analysis of the RRGS data using PCA and Admixture identified significant geographical structure to genetic variation, but patterns of population structure were almost identical in each subgenome. The PCA evidenced genetic differentiation within *X. laevis*, with the first and second principal component distinguishing from the summer, winter, northwest transitional rainfall zone, and the southeast transitional rainfall zone (Fig. 3), even though these components accounted for a small proportion of the total genotypic variation (∼7%). Samples from Nieuwoudtville in the northwest transition zone are more distinguished from populations in the summer or winter rainfall zones than are samples from the southeast transition zone from populations in the summer or winter rainfall zones.

**Fig. 2.**
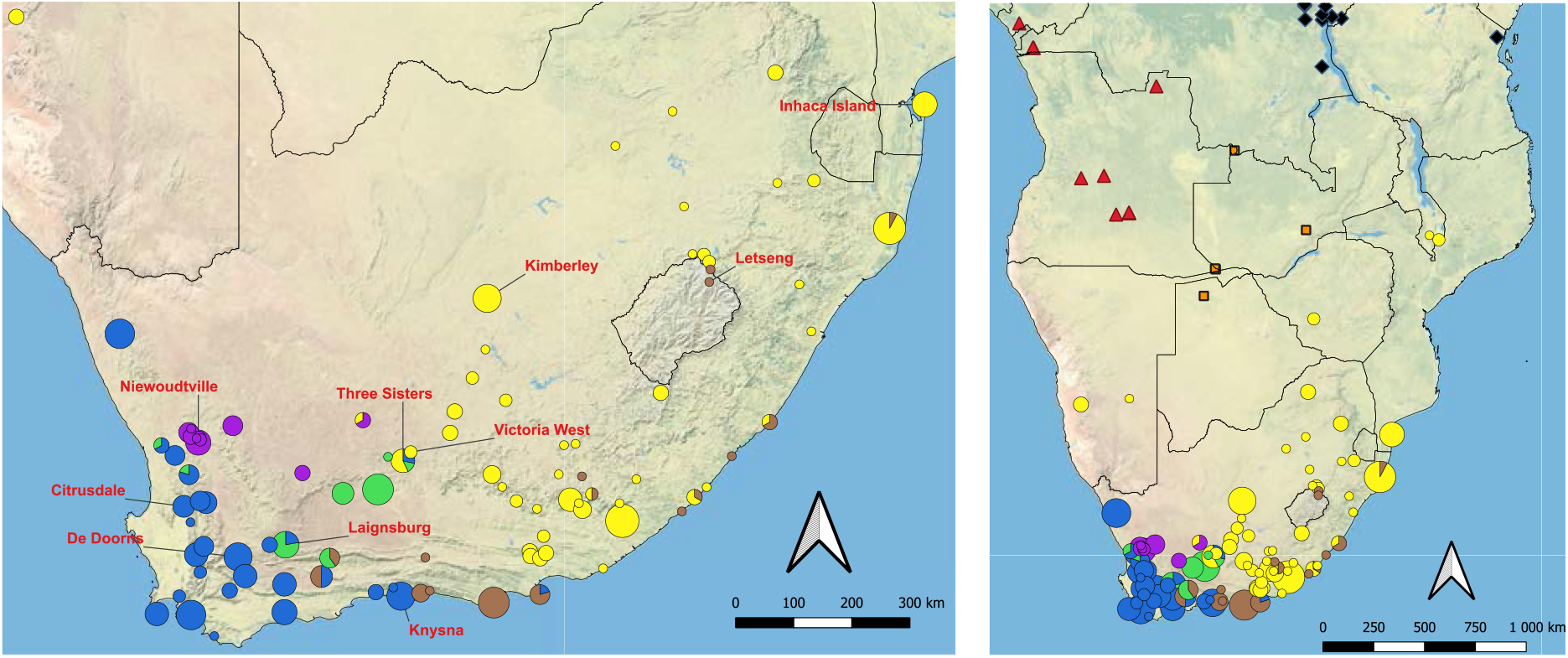
Geographical distribution of *X. laevis* sampling localities in South Africa (including Lesotho) used for RRGS (red labels, left panel) and Sanger sequencing of mitochondria (circular pies, triangles, squares, and diamonds for *X. laevis, X. petersii, X. poweri*, and *X. victorianus* respectively). For *X. laevis*, colors correspond to mitochondrial clades in Fig. 5. In the left panel, solid and dotted lines indicate the margins of the winter rainfall zone to the southwest and the summer rainfall to the northeast, respectively (Chase and Meadows 2007). The locality of the *X. gilli* sample, not shown, falls within the range of the winter rainfall (blue) clade of *X. laevis*, and is listed in Table S2.

**Fig. 3.**
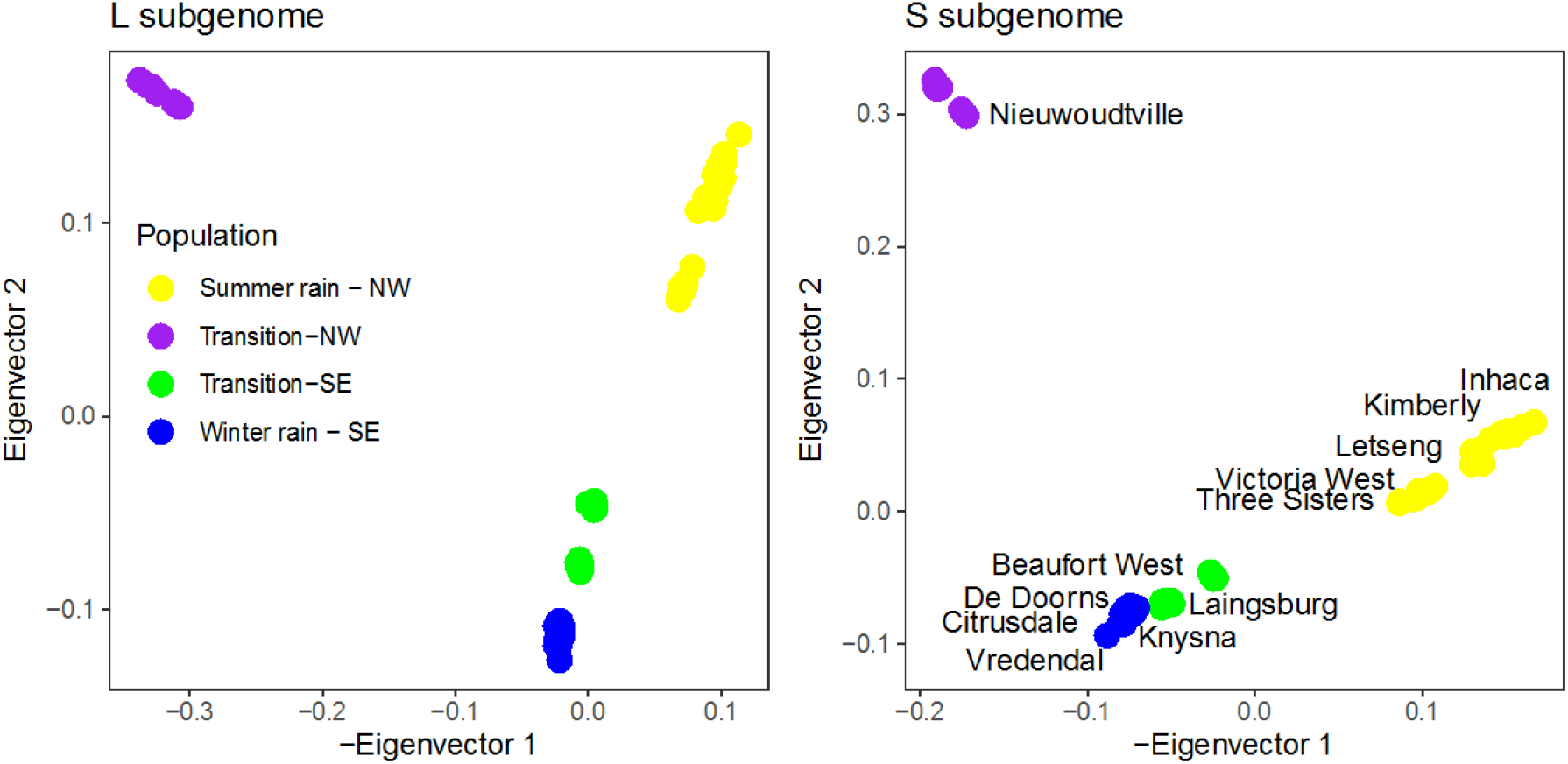
Principal components analysis of RRGS data from *X. laevis* for the L and the S subgenomes (right and left, respectively). For the L subgenome, the first and second eigenvectors explain 3.5% and 3.3% of genotypic variation respectively; For the S subgenome, the first and second eigenvectors explain 3.6% and 3.4% of genotypic variation, respectively. Patterns of population structure are similar in each subgenome and samples cluster by rainfall zone (indicated with color in the left panel) and by locality (labeled in right panel).

Admixture analysis recovered the most prominent structure between populations on either side of the southwest margin of the Great Escarpment. This is apparent when *k* = 2 where samples to the southwest of this feature (Knysna, De Doorns, Citrusdale, Vredendal) carry one ancestry component and those to the northwest (Beaufort West, Three Sisters, Victoria West, Kimberly, Letseng, Inhaca) carry another, with admixed ancestry components in Laingsburg and Nieuwoudtville (Fig. 4). At higher values of *k* additional population structure is evident, and patterns of population subdivision are similar to the PCA analysis. For example, at *k* = 5, animals from Three Sisters are distinguished from their neighbors to the north (Victoria West, Kimberly, Letseng, Inhaca) and those to the south (Knysna, De Doorns, Citrusdale, Laingsburg, Beaufort West).

**Fig. 4.**
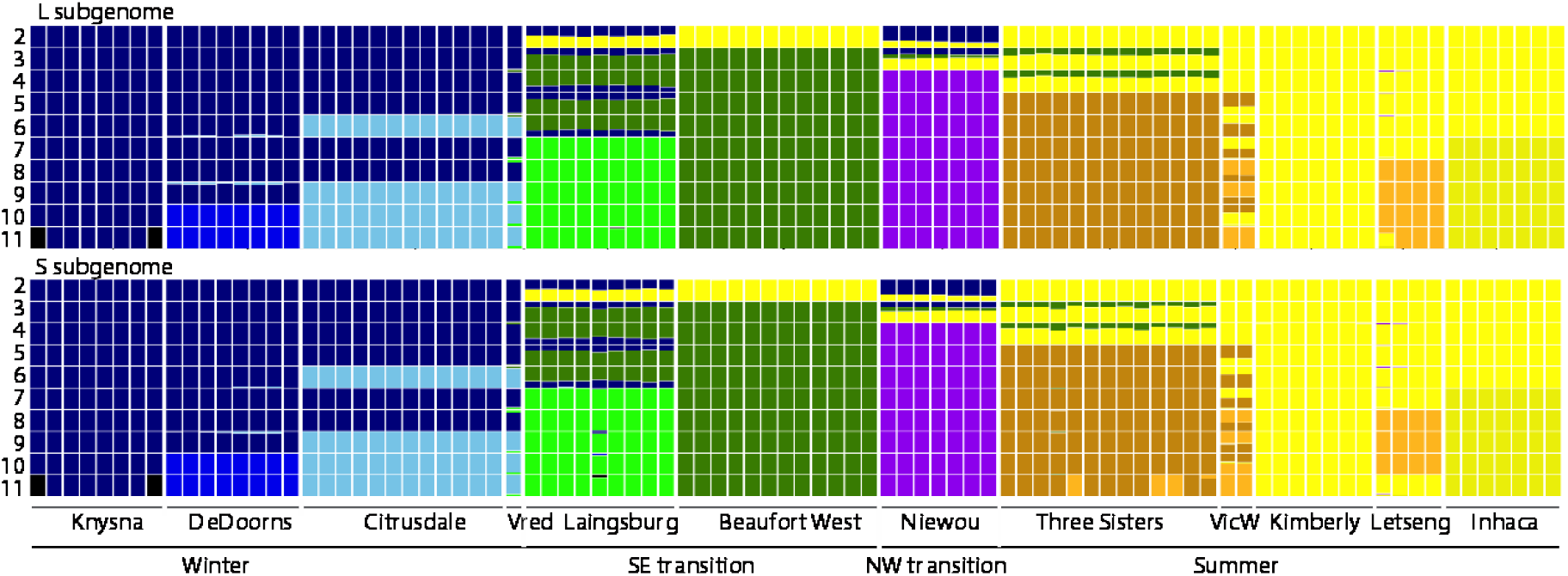
Admixture plot using only the L subgenome (top) or only the S subgenome (bottom) for 2-11 ancestry components (*k*) labeled on the left side. The order of the samples is from left to right matches the order of samples in Table S1.

These analyses also provide resolution into areas of admixture, such as *X. laevis* from Laingsburg, where most ancestry components are shared with individuals from Beaufort West which lies to the northeast of Laingsburg, but where individuals also carry an ancestry component that is most common in populations in the winter rainfall zone to the southwest. Similarly, *X. laevis* from Victoria West have an ancestry shared with *X. laevis* populations on the top of the Great Escarpment and beyond to Mozambique (Kimberly, Letseng, Inhaca) but also carry ancestry components from Three Sisters, which is situated on the approach to the escarpment. Based on the cross-validation procedure of Admixture (Alexander et al. 2009), the best value of *k* for these data is 5 or 6 for each subgenome (Supplementary Table 3).

### No evidence for differential geneflow by subgenome

We performed four non-independent analyses of introgression based on Patterson’s *D* statistic using four combinations of four populations. For three of these four analyses (Comparisons 1, 3, and 4), Patterson’s *D* was significantly greater than zero but not significantly different for each subgenome based on the permutation tests (Table 1). The weighted average of Patterson’s *D* was more extreme (but not significantly so) in the S than the L subgenome for Comparison 2. Overall, this is consistent with extensive, geographically structured gene flow, but does not provide evidence for differential gene flow by subgenome in most comparisons on a genome-wide scale. For all four comparisons, the effect size is below 0.2, suggesting that the effect of subgenome on Patterson’s *D* is weak (Table 1). The test statistics are generally far smaller than the value estimated to have 80% power to detect a significant difference. This indicates that the data are underpowered to detect very weak effects of subgenome on Patterson’s *D*, if one exists. When only genic areas were considered, results were similar to the analysis of all RRGS data within each subgenome, with three of four comparisons having more extreme values of Patterson’s *D* statistic in the L rather than the S subgenome, and with the values in Comparison 2 being not significantly different (Table S4).

**Table 1.**
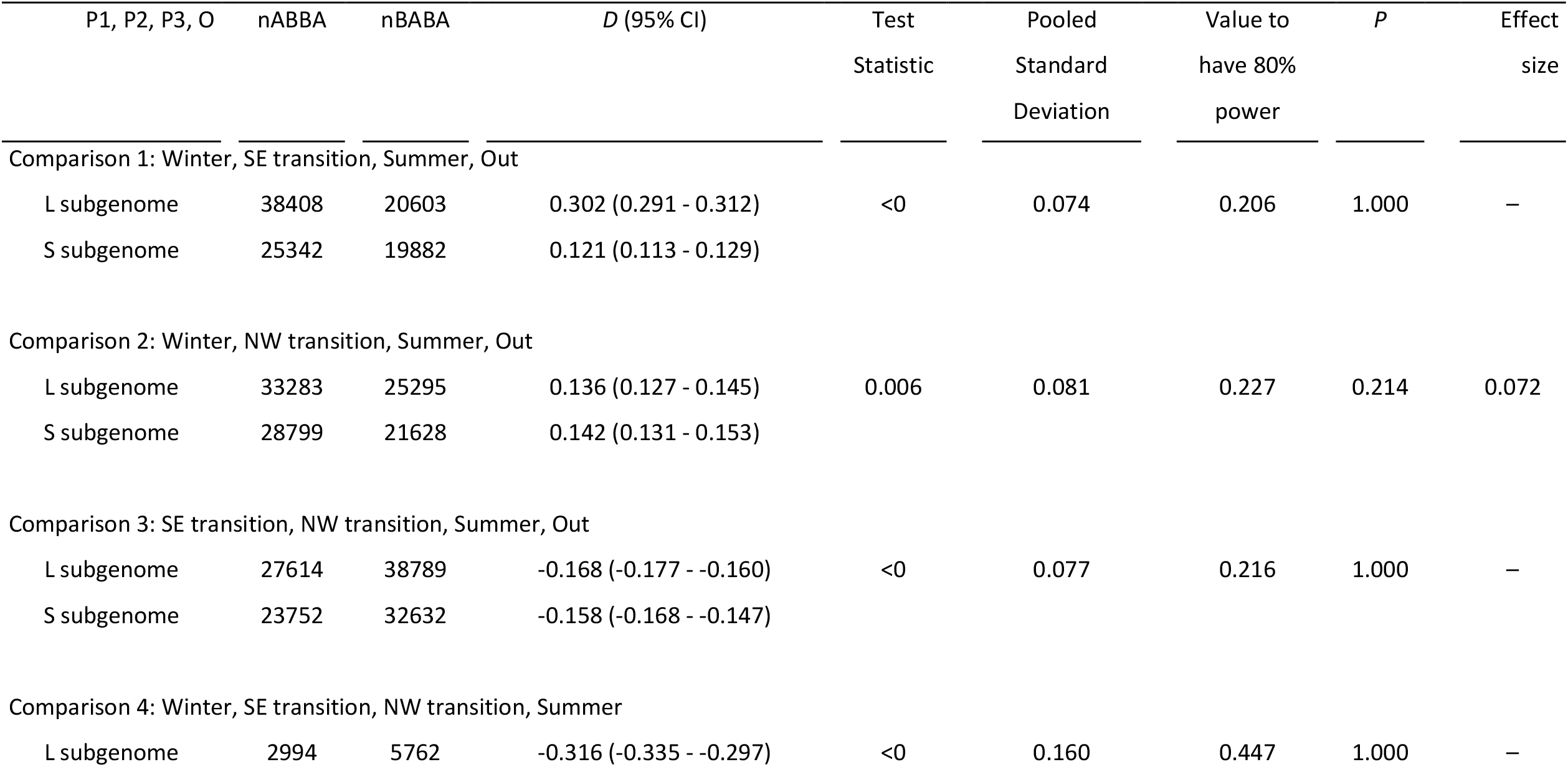
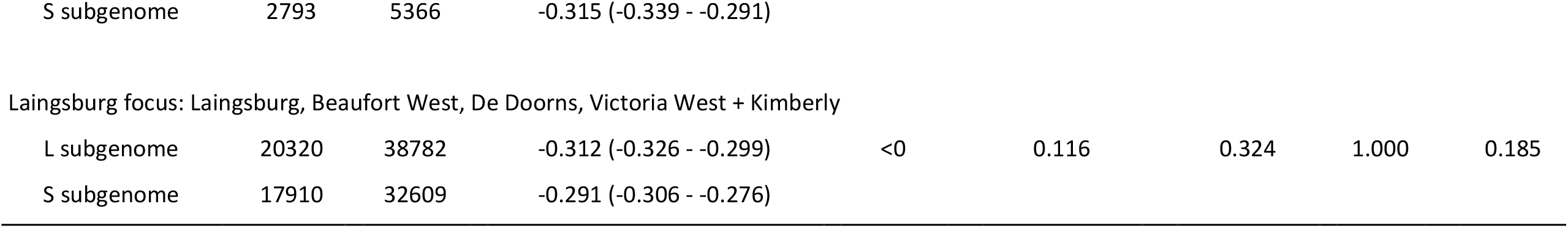
Patterson’s *D* statistic and 95% confidence intervals (95% CI) for each subgenome for four comparisons within *X. laevis* involving populations grouped by rainfall pattern (Winter, NW transition, SE transition, Summer, and the *X. victorianus* + *X. poweri* outgroup (Out) and a focused analysis of the zone of admixture in Laingsburg. P values of permutation tests (P) indicate that the *D* statistic is not significantly different between the L and the S subgenome for any comparison. For each comparison and for each subgenome the number of ABBA and BABA sites (nABBA, nBABA, defined in main text) is indicated. For each comparison the test statistic, pooled standard deviation, value of the test statistic that would deliver 80% power, P value, and effect size is also provided. The effect size of relationships with a test statistic below zero are not shown.

| P1, P2, P3, O | nABBA | nBABA | *D* (95% CI) | Test<br>Statistic | Pooled<br>Standard<br>Deviation | Value to<br>have 80%<br>power | *P* | Effect<br>size |
| --- | --- | --- | --- | --- | --- | --- | --- | --- |
| Comparison 1: Winter, SE transition, Summer, Out |  |  |  |  |  |  |  |  |
| L subgenome | 38408 | 20603 | 0.302 (0.291 - 0.312) | <0 | 0.074 | 0.206 | 1.000 | – |
| S subgenome | 25342 | 19882 | 0.121 (0.113 - 0.129) |  |  |  |  |  |
| Comparison 2: Winter, NW transition, Summer, Out |  |  |  |  |  |  |  |  |
| L subgenome | 33283 | 25295 | 0.136 (0.127 - 0.145) | 0.006 | 0.081 | 0.227 | 0.214 | 0.072 |
| S subgenome | 28799 | 21628 | 0.142 (0.131 - 0.153) |  |  |  |  |  |
| Comparison 3: SE transition, NW transition, Summer, Out |  |  |  |  |  |  |  |  |
| L subgenome | 27614 | 38789 | -0.168 (-0.177 - -0.160) | <0 | 0.077 | 0.216 | 1.000 | – |
| S subgenome | 23752 | 32632 | -0.158 (-0.168 - -0.147) |  |  |  |  |  |
| Comparison 4: Winter, SE transition, NW transition, Summer |  |  |  |  |  |  |  |  |
| L subgenome | 2994 | 5762 | -0.316 (-0.335 - -0.297) | <0 | 0.160 | 0.447 | 1.000 | – |

|  |  |  |  |
| --- | --- | --- | --- |
| S subgenome | 2793 | 5366 | -0.315 (-0.339 - -0.291) |
| Laingsburg focus: Laingsburg, Beaufort West, De Doorns, Victoria West + Kimberly |  |  |  |  |  |  |  |  |
| L subgenome | 20320 | 38782 | -0.312 (-0.326 - -0.299) | <0 | 0.116 | 0.324 | 1.000 | 0.185 |
| S subgenome | 17910 | 32609 | -0.291 (-0.306 - -0.276) |  |  |  |  |  |

Table 1. Patterson's *D* statistic and 95% confidence intervals (95% CI) for each subgenome for four comparisons v (Winter, NW transition, SE transition, Summer, and the *X.
| P1, P2, P3, O | nABBA | nBABA | D (95% CI) | Test Statistic |
| --- | --- | --- | --- | --- |
| Comparison 1: Winter, SE transition, Summer, Out |  |  |  |  |
| L subgenome | 38408 | 20603 | 0.302 (0.291 - 0.312) | <0 |
| S subgenome | 25342 | 19882 | 0.121 (0.113 - 0.129) |  |
| Comparison 2: Winter, NW transition, Summer, Out |  |  |  |  |
| L subgenome | 33283 | 25295 | 0.136 (0.127 - 0.145) | 0.006 |
| S subgenome | 28799 | 21628 | 0.142 (0.131 - 0.153) |  |
| Comparison 3: SE transition, NW transition, Summer, Out |  |  |  |  |
| L subgenome | 27614 | 38789 | -0.168 (-0.177 - -0.160) | <0 |
| S subgenome | 23752 | 32632 | -0.158 (-0.168 - -0.147) |  |
| Comparison 4: Winter, SE transition, NW transition, Summer |  |  |  |  |
| L subgenome | 2994 | 5762 | -0.316 (-0.335 - -0.297) | <0 |
| S subgenome | 2793 | 5366 | -0.315 (-0.339 - -0.291) |  |
| Laingsburg focus: Laingsburg, Beaufort West, De Doorns, Victoria West + Kimberly |  |  |  |  |
| L subgenome | 20320 | 38782 | -0.312 (-0.326 - -0.299) | <0 |
| S subgenome | 17910 | 32609 | -0.291 (-0.306 - -0.276) |  |

| <u>Pooled</u><br><u>Standard</u><br><u>Deviation</u> | <u>Value to</u><br><u>have 80%</u><br><u>power</u> | <u>*P*</u> | <u>Effect size</u> |
| --- | --- | --- | --- |
| 0.074 | 0.206 | 1.000 | - |
| 0.081 | 0.227 | 0.214 | 0.072 |
| 0.077 | 0.216 | 1.000 | - |
| 0.160 | 0.447 | 1.000 | - |
| 0.116 | 0.324 | 1.000 | - |

For Comparison 1, Patterson’s *D* is positive for both subgenomes, supporting the geographically based intuition that there is more gene flow between summer rainfall and the southeast (SE) transition than between summer rainfall and winter rainfall. Similarly, for Comparison 2 Patterson’s *D* is also positive, indicating more gene flow between summer rainfall and the northwest (NW) transition than between summer rainfall and winter rainfall (which is also unsurprising since the NW transition is closer than the winter rainfall region to the summer rainfall region). For Comparison 3, Patterson’s *D* is negative, indicating more gene flow between summer rainfall and the SE transition than between summer rainfall and NW transition. This result is consistent with the PCA and admixture analyses discussed above, which highlight the distinctiveness of the NW transition population. For Comparison 4, Patterson’s *D* is also negative, indicating more geneflow between NW transition and winter rainfall than between NW transition and SE transition. This suggests that gene flow within the transition zone is more limited than between the transition zone and the neighboring regions, a finding that is further highlighted by generally non-overlapping mitochondrial clades as detailed below.

Isolation by distance is expected to differently affect population structure within the summer, winter, and NW and SE transition zones due to differences in the size of each region. For this reason, we performed an additional analysis focused on the admixed population of Laingsburg as a way of exploring whether there might be subgenome effects at a smaller geographic scale (Laingsburg focus, Table 1). Contrary to this possibility but largely consistent with findings from Comparisons 1 – 4, this analysis found a more extreme Patterson’s *D* statistic in the L subgenome than in the S subgenome (Table 1). For both subgenomes, Patterson’s *D* was negative for the Laingsburg focus comparison, which indicates more genetic exchange between the geographically proximal populations in De Doorns and Laingsburg as compared to the more distant comparison between De Doorns and Beaufort West. When only genic regions were considered, there also was a more extreme (negative) Patterson’s *D* statistic in the L rather than the S subgenome (Table S4).

Subgenome-wide genetic similarity as summarized by phenograms also suggested a high degree of congruence between the L and S subgenomes (Fig. 5). Phenetic relationships among nuclear variation are similar in each subgenome and indicate higher similarity within *X. laevis* individuals compared to between *X. laevis* and either *X. victorianus* or *X. poweri* (Fig. 5). Within *X. laevis*, similarity within each population (winter rainfall, southeast transition, northwest transition, summer rainfall) is higher than between these populations (Fig. 5).

**Fig. 5.**
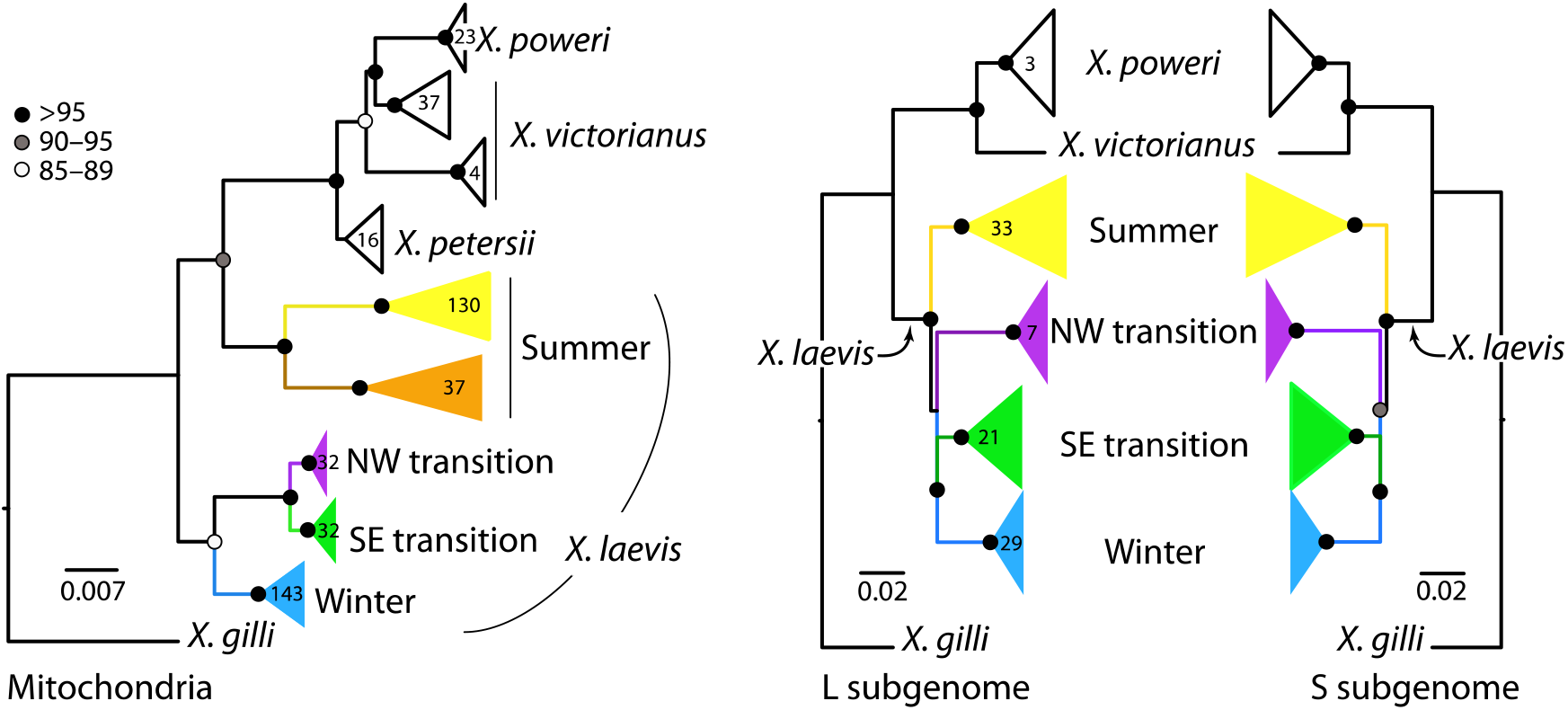
Evolutionary relationships among mitochondrial sequences (left) are different from similarity in the L and S subgenomes (center and right, respectively). Shaded nodes indicate bootstrap values as indicated on the scale; numbers inside clades indicate the sample sizes of individuals; samples sizes of the S subgenome (not shown) are identical to those for the L subgenome. The scale bar indicates substitutions per site for the mitochondrial phylogeny; branch lengths of the phenogram are scaled by only variable positions and are not comparable to mitochondrial branch lengths. Samples in each clade of the mitochondrial tree are listed in Supplementary Table 1. Samples with the brown mitochondrial lineage (left) were not present in the RRGS data (center, right).

Because population dynamics of mitochondrial DNA may not be fully captured by phylogenetic analyses that assume strictly bifurcating relationships, we also constructed a network of relationships among the partial mitochondrial sequences (Fig. 6). This network identified one star-shaped relationships in the winter “blue” clade and two in the summer “yellow” clade that are suggestive of population expansion (Ferreri et al. 2011). Several haplotype clusters in the network are separated by multiple steps, which is consistent with divergence between the clusters. The brown clade has comparatively more missing/inferred haplotypes than the blue and green clades, which suggests a more stable population size in the brown clade. Relatively few changes separate the green and purple clades, which is consistent with the close inferred phylogenetic relationship between these clades (Fig. 5).

**Fig. 6.**
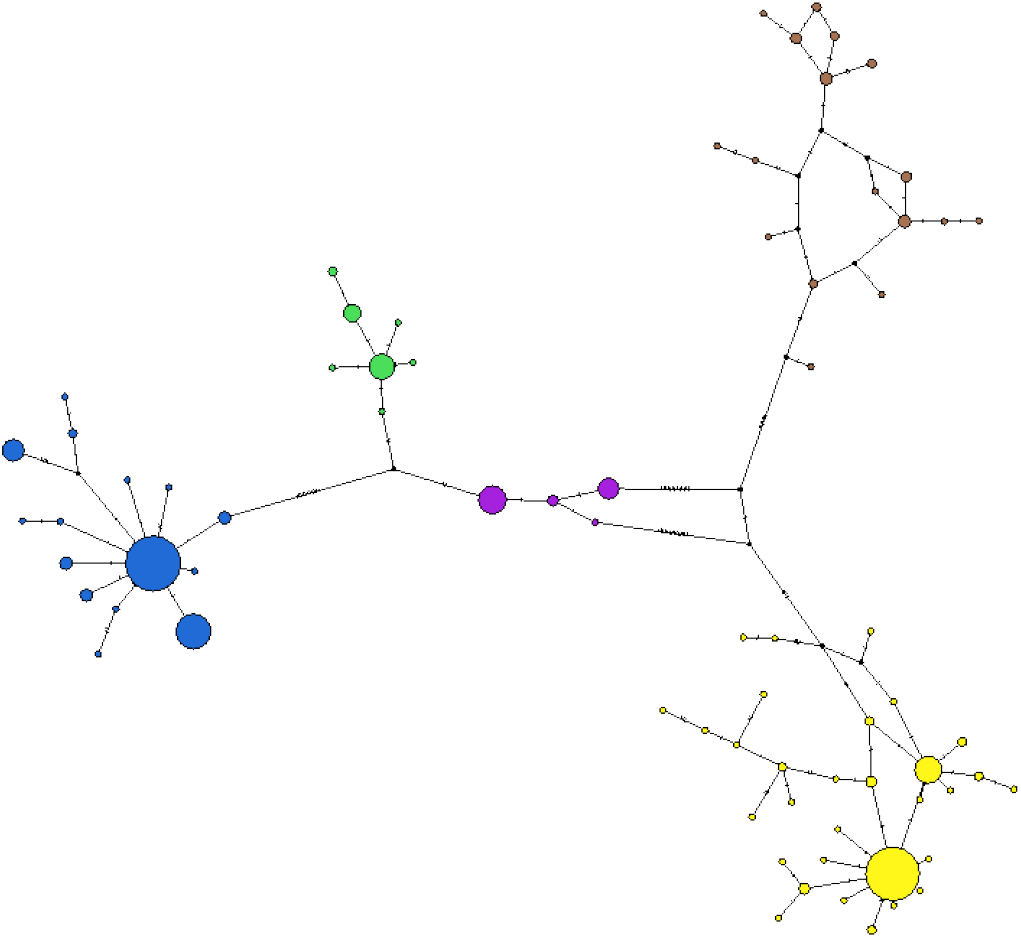
Parsimony network of partial *X. laevis* mitochondrial sequences. Colors correspond to Fig. 2; the size of the nodes is proportional to the number of samples with each haplotype. Inferred nodes are indicated with black nodes; hashes on branches indicate changes between nodes that are not represented by a sampled sequence.

### Phylogeography of X. laevis in South Africa

Major topographic relationships recovered from the phylogenetic and phenetic analysis (Fig. 5) are consistent with previous analyses of fewer partial mitochondrial sequences or concatenated nuclear loci (Furman et al. 2015). These analyses suggest that mitochondrial DNA within *X. laevis* is paraphyletic relative to *X. victorianus, X. poweri*, and *X. petersii*, a result that is discordant with phenetic analysis of nuclear DNA which finds lower intraspecific than interspecific divergence in both subgenomes (discussed above). Mitochondrial sequences support a sister relationship between the northwest and southeast transition zones; this relationship was unresolved in the phenetic analysis of each subgenome (Fig. 5).

Increased sampling identified a previously unreported mitochondrial clade (Fig. 3; hereafter “brown”) that is sister to the winter rainfall (yellow) clade. The brown clade is prevalent in the southeastern portion of the distribution of *X. laevis* east of the Keurbooms River and is also found at lower frequencies in animals that were sampled north of this area in the Letseng, where the yellow clade is common. The winter rainfall clade covers most of the range of *X. laevis* in southern Africa, including populations sampled in Namibia, Zimbabwe, Malawi, and Mozambique.

New sampling of mitochondrial sequences also permitted us to identify the geographic margins of previously identified mitochondrial clades. For example, the northwest transition zone (purple) clade, which was previously known only from Nieuwoudtville at the northern extent of the Cape Fold Mountains (Furman et al. 2015), extends east onto the Great Escarpment in the Karoo (Fig. 1). The southeast transition zone (green) clade, which was previously known from Laingsburg and Beaufort West, also occurs on the inner margins of the Cape Fold Mountains (north-south and east-west), mainly within the Karoo basin below the Great Escarpment. The Winter rainfall (blue) clade is carried by animals from the southwestern extreme of the distribution of *X. laevis* and at least as far north as Springbok, and with the southeast margin near Knysna.

Multiple mitochondrial clades were observed at the following localities: Vredendal (blue, green), Three Sisters (blue, green, yellow), Port Elizabeth (blue, brown), Roiiberg Lodge (blue, brown), Calitzdorp Dam (Brown, Green), Manubi Forest (brown, yellow), Katberg (brown, yellow), Alpha Excelsior Farm (blue, green), Mkambati Horseshoe Falls (brown, yellow), Malala Lodge (brown, yellow). This geographic comingling of mitochondrial clades at the margins of their distributions corresponds with evidence multiple ancestry components in autosomal DNA from the same individuals from at least two localities (Vredendal, Three Sisters).

## Discussion

### Absence of a substantial subgenome effect for most comparisons

Subgenome evolution is not universally asymmetric (e.g., Liu et al. 2017; Wu et al. 2021) and the effect of subgenome on various genomic characteristics (gene flow, gene expression, etc.) may be modest, species-specific, temporally dynamic, or centered only on small genomic regions.

Our analyses of genetic similarity, population structure, and introgression failed to detect significant or substantial differences between the L and S subgenomes in natural populations of *X. laevis* in multiple comparisons with all the RRGS data, and in multiple comparisons with RRGS data from genic regions. In fact, for four of five comparisons in each category (all RRGS data or genic regions only), Patterson’s *D* statistic was more extreme (positive or negative) for the L subgenome than the S subgenome; for the fifth comparison (Comparison 2 in both cases) there was no significant difference between the L and S subgenomes. Overall, these findings are inconsistent with the possibility that the L subgenome plays a disproportionately large role in local adaptation or that gene flow in the S subgenome is atypically high in order to mitigate deleterious effects of the polyploid ratchet.

Subgenomes do have significant and substantial effects on several other aspects of *X. laevis* genome evolution, including transposable element mobility and composition, genomic rearrangements, gene silencing, expression, and length, and the strength of purifying selection on genic regions (Session et al. 2016; Elurbe et al. 2017; Furman et al. 2018). One possibility is that differential introgression within subgenomes could occur in small genomic regions that we lacked statistical power to discern. An interesting question that could be explored with additional high quality genome assemblies from other allotetraploid *Xenopus* species (in subgenus *Xenopus*) asks whether and how the rate of genomic rearrangements has varied over time since allotetraploidization. One possibility is that purifying selection counteracted the polyploid ratchet by eliminating rearrangements that occurred recently. If this were the case, many rearrangements in the S subgenome of *X. laevis* (Session et al. 2016) would be shared with other allotetraploid species that diversified soon after allopolyploidization.

### Genetic differentiation of the nuclear genome of X. laevis

Previously, patterns of population structure within the natural range of *X. laevis* were inferred from genetic variation in portions of the mitochondrial genome and 2–15 nuclear genes (Grohovaz et al. 1996; Evans et al. 1997; Measey and Channing 2003; Du Preez et al. 2009; Furman et al. 2015).

These efforts identified significant intraspecific differentiation, and admixture in a population from Laingsburg from source populations to the southwest and northeast. The far larger sample of genetic variation afforded by RRGS data examined here further contextualizes the geographic distribution of genetic variation in *X. laevis*, including the distribution of ancestry components at two localities (Citrusdale and Three Sisters) that have not previously been studied.

Our results are consistent with Furman et al. (2015) in identifying substantial population structure in *X. laevis* from South Africa (including Lesotho), particularly between individuals sampled near Nieuwoudtville, which is the northwest transition zone on top of the Great Escarpment. The distribution of differentiated populations roughly matches the distribution of mitochondrial clades (purple, yellow, blue + green respectively). In the RRGS data there were no samples that carried the brown clade, so we were unable to assess genetic variation in their nuclear genomes. Patterson’s *D* provides geographically plausible inferences of gene flow between populations, including higher levels of gene flow between geographically adjacent compared to non-adjacent populations. This statistic also highlights the population isolation of *X. laevis* from Nieuwoudtville: gene flow between summer rainfall region and the SE transition zone is higher than between summer rainfall region and the NW (Nieuwoudtville) transition zone (evidenced by a negative *D* in comparison 3, Table 1). Interestingly, gene flow within the transition zone is modest: introgression between the NW transition zone and the winter rainfall zone is higher than between the NW transition zone and the SE transition zone (evidenced by a negative *D* in comparison 4, Table 1).

Analysis of genetic variation in the nuclear genome further characterized several zones of admixture, including one that was previously identified (Laingsburg; Du Preez et al. 2009), and one that was previously studied with less information and whose ancestry components were incompletely characterized (Victoria West; Furman et al. 2015). In Laingsburg, ancestry of individuals was primarily shared with Beaufort West (which lies to the northeast of Laingsburg) but also with a component from the winter rainfall zone to the southwest of Laingsburg. Previously uncharacterized genetic variation from Three Sisters, which lies between Beaufort West and Victoria West, highlights a gradient of ancestry components as one moves through the transitional zone (Fig. 2) from southwest to northeast towards the top of the Great Escarpment. New information from Citrusdale and Vredendal confirms that the winter rainfall (“blue”) population occurs throughout this region.

New samples analyzed here from Sentinel Peak in Letseng, a high altitude area in the Drakensburg portion of the Great Escarpment, which is >3,000 m above sea level (asl), were not substantially differentiated from a population near Kimberly, which is ∼1,100 m asl. This suggests that genome-wide genetic changes associated with adaptation to high altitude (see Wagener et al. 2021) are small in comparison to population structure within *X. laevis*.

### Mitochondrial variation; margins of clades

The first phylogeographic exploration of mitochondrial variation in *X. laevis* showed the existence of the winter rainfall clade (blue), the summer rainfall clade (yellow), and an isolated sample from the NW transition zone – Nieuwoudtville (purple) as a divergent clade (Grohovaz et al. 1996).

Subsequent phylogeographic studies also found the NW transition clade in and around Nieuwoudtville (Measey and Channing 2003; Furman et al. 2015). Here we show that this clade is also carried in animals east of Nieuwoudtville onto the great escarpment and into the Karoo.

Nieuwoudtville is thus at the western edge of the distribution of this clade, with animals from the winter rainfall clade being found at similar altitude immediately south, and below the escarpment to the west. Our sampling was unable to characterize the northern extent of the NW transition clade, as this area is especially arid and contains the least records in the region for this species (Measey 2004b).

Inferred relationships between the mitochondrial sequences differ in several ways from patterns of genome-wide similarity in the nuclear genome, including within *X. laevis* and between *X. laevis* and other closely related species. Ancestral polymorphism is a plausible explanation for these discrepancies. Male-biased migration contributes to mito-nuclear phylogenomic discordance in some species, but there is no evidence for this in *X. laevis* (Measey 2016; Courant et al. 2017; De Villiers and Measey 2017; Louppe et al. 2020).

The distribution of the mitochondrial clade from the southeast transition zone (green), first identified by Du Preez et al. (2009) in Beaufort West, was demonstrated by this study to also occupy most of the Karoo basin below the Great Escarpment, including the inner montane areas to the west and south. The distribution of the winter rainfall clade (blue) is shown to occur mostly in lowland areas and extending north at least as far as Springbok, and into the Cape Fold Mountains.

The winter rainfall clade also extends into low lying regions in the southern portion of the transition zone, with the eastern margin at the Keurbooms River. Animals sampled east of this river carried a previously unrecognized mitochondrial clade (brown in Fig. 1 and 2), including as far north as the Drakensburg mountains, or alternatively the widespread summer rainfall (yellow) clade. The summer rainfall clade encompasses all populations of *X. laevis* away from this southwestern corner of southern Africa, including Namibia, Zimbabwe, Mozambique, and Malawi. Measey (2004a) showed that this is the most widely distributed of all amphibian species in South Africa, with records from nearly every quarter degree square. Areas for further investigation include (a) the limit of the winter rainfall (blue) clade in the far Northern Cape, which may be the Fish River at the border with Namibia based on findings in other amphibians (see Tolley *et al*. 2014), (b) the northern extent of the purple clade which could be bounded by the arid regions toward the Kgalagadi Transfrontier Park that straddles the border between South Africa and Botswana near Namibia, and (c) the eastern limit of the green clade, which may extend into the eastern Karoo region.

## Conclusions

Genetic variation from the nuclear genome of an allopolyploid frog – *Xenopus laevis* – in their natural habitat failed to identify substantial differences in population structure or gene flow in each subgenome of this allotetraploid species. These results are inconsistent with the proposal that a polyploid ratchet favors restorative gene flow in the S subgenome – at least on a scale that is substantial enough to detect using the RRGS data we collected. This suggests that the effect of subgenomes on gene flow and population structure is either small or absent across the genome of *X. laevis*. Population structure in the nuclear genome includes several zones of admixture, and corresponds roughly with distributions of mitochondrial clades, ecological and rainfall transition zones, and topographic relief associated with the Cape Fold Mountains, although surprisingly less so for the Drakensburg Mountains.

## Acknowledgements

This research was supported by the Natural Science and Engineering Research Council of Canada (RGPIN-2017-05770), a Resource Allocation Competition awards from Compute Canada, the Museum of Comparative Zoology at Harvard University, and The National Research Foundation (NRF) of South Africa (NRF Grant No. 87759 to JM) provided financial support for the project sampling and sequencing. JM was supported by the DSI-NRF Centre of Excellence for Invasion Biology. Fieldwork by WC was supported by the Foundational Biodiversity Information Program, BioGaps, Bayworld, and the Eastern Cape Parks and Tourism Agency. We thank CapeNature, the Eastern Cape Department of Economic Development, Environmental Affairs and Tourism; Eastern Cape Parks and Tourism Agency; Angolan Ministry of Environment Institute of Biodiversity (MINAMB) for permits. We would like to thank to following individuals for donating tissues: Andrew Turner, Sarah Davies, Jaco le Roux, Likho Sikutshwa, Mohlamotsane Mokhatla, Nick Telford, Ross Shackleton, Alex Rebelo, Olaf Weyl, Ana Nunes, Andre de Villiers, Louis du Preez, Retha Hofmeyr, Ben Smit and Rob Hopkins. We would also like to thank Krystal Tolley (SANBI) and Roger Bills (SAIAB) for providing subsamples of tissues in their collections and the property owners of localities listed in Table S2.

## References

Alexander, D. H., J. Novembre, and K. Lange. 2009. Fast model-based estimation of ancestry in unrelated individuals. Genome Research 19:1655–1664.

Bewick, A. J., D. W. Anderson, and B. J. Evans. 2011. Evolution of the closely related, sex-related genes DM-W and DMRT1 in African clawed frogs (Xenopus). Evolution 65:698–712.

Bolger, A. M., M. Lohse, and B. Usadel. 2014. Trimmomatic: a flexible trimmer for Illumina sequence data. Bioinformatics 30:2114–2120.

Bomblies, K. 2020. When everything changes at once: finding a new normal after genome duplication. Proceedings of the Royal Society Biological Sciences Series B 287:Article No.: 20202154.

Busing, F. M. T. A., E. Meijer, and R. van der Leeden. 1999. Delete-m jackknife for unequal m. Statistics and Computing 9:3–8.

Cannatella, D. C. and R. O. de Sá. 1993. Xenopus laevis as a model organism. Systematic Biology 42:476–507.

Chase, B. M. and M. E. Meadows. 2007. Late Quaternary dynamics of southern Africa’s winter rainfall zone. Earth-Science Reviews 84:103–138.

Cheng, F., J. Wu, X. Cai, J. Liang, M. Freeling, and X. Wang. 2018. Gene retention, fractionation and subgenome differences in polyploid plants. Nature Plants 4:258–268.

Clement, M., D. Posada, and K. A. Crandall. 2000. TCS: a computer program to estimate gene genealogies. Molecular Ecology 9:257–270.

Cohen, J. 2013. Statistical Power Analysis for the Behavioral Sciences. Routledge Academic, New York.

Courant, J., S. Vogt, R. Marques, J. Measey, J. Secondi, R. Rebelo, A. De Villiers, F. Ihlow, C. De Busschere, T. Backeljau, D. Roedder, and A. Herrel. 2017. Are invasive populations characterized by a broader diet than native populations? PeerJ 5: e3250.

Daudin, F. M. 1802. Histoire Naturelle des Rainettes, des Grenouilles et des Crapauds. Levrault, Paris.

De Busschere, C., J. Courant, A. Herrel, R. Rebelo, R. Rödder, G. J. Measey, and T. Backeljau. 2016. Unequal contribution of native South African phylogeographic lineages to the invasion of the African clawed frog, Xenopus laevis, in Europe. PeerJ 4:e1659

De Villiers, F. A. and J. Measey. 2017. Overland movement in African clawed frogs (Xenopus laevis): empirical dispersal data from within their native range. PeerJ 5: e4039.

Development_team. 2019. Picard Toolkit. Broad Institute, GitHub Repository. https://broadinstitute.github.io/picard/.

Du Preez, L. H. and V. C. Carruthers. 2017. Frogs of southern Africa. Penguin Random House, Cape Town, South Africa.

Du Preez, L. H., N. Kunene, R. Hanner, J. P. Giesy, K. R. Solomon, A. Hosmer, and G. Van Der Kraak. 2009. Population-specific occurrence of testicular ovarian follicles in Xenopus laevis from South Africa. Aquatic Toxicology 95:10–16.

Edger, P. P., T. Poorten, R. VanBuren, M. A. Hardigan, M. Colle, M. R. McKain, R. D. Smith, S. Teresi, A. D. L. Nelson, C. M. Wai, E. Alger, I K. A. Bird, A. E. Yocca, N. Pumplin, S. Ou, G. Ben-Zvi, A. Brode, K. Baruch, T. Swale, L. Shiue, C. B. Acharya, G. S. Cole, J. P. Mower, K. L. Childs, N. Jiang, E. Lyons, M. Freeling, J. R. Puzey, and S. J. Knapp. 2019. Origin and evolution of the octoploid strawberry genome. Nature Genetics 51:541–547.

Elurbe, D. M., S. S. Paranjpe, G. Georgiou, I. Van Kruijsbergen, O. Bogdanovic, R. Gibeaux, R. Heald, R. Lister, M. A. Huynen, S. J. Van Heeringen, and G. J. C. Veenstra. 2017. Regulatory remodeling in the allo-tetraploid frog Xenopus laevis. Genome Biology 18:Article No.: 198.

Evans, B. J. 2008. Genome evolution and speciation genetics of allopolyploid clawed frogs (Xenopus and Silurana). Frontiers in Bioscience 13:4687–4706.

Evans, B. J., R. M. Brown, J. A. McGuire, J. Supriatna, N. Andayani, A. Diesmos, D. T. Iskandar, D. J. Melnick, and D. C. Cannatella. 2003. Phylogenetics of Fanged Frogs (Anura; Ranidae; Limnonectes): testing biogeographical hypotheses at the Asian-Australian faunal zone interface. Systematic Biology 52:794–819.

Evans, B. J., T. F. Carter, E. Greenbaum, V. Gvoždík, D. B. Kelley, P. J. McLaughlin, O. S. G. Pauwels, D. M. Portik, E. L. Stanley, R. C. Tinsley, M. L. Tobias, and D. C. Blackburn. 2015. Genetics, morphology, adverstisement calls, and historical records distinguish six new polyploid species of African clawed frog (Xenopus, Pipidae) from West and Central Africa. PLoS One 10:e0142823 (0142851 pages).

Evans, B. J., M. T. Gansauge, E. L. Stanley, B. L. S. Furman, C. M. S. Cauret, C. Ofori-Boateng, V. Gvozdik, J. W. Streicher, E. Greenbaum, R. C. Tinsley, M. Meyer, and D. C. Blackburn. 2019. Xenopus fraseri: Mr. Fraser, where did your frog come from? PLoS One 14:e0220892 (0220814 pages).

Evans, B. J., D. B. Kelley, R. C. Tinsley, D. J. Melnick, and D. C. Cannatella. 2004. A mitochondrial DNA phylogeny of clawed frogs: Phylogeography on sub-Saharan Africa and implications for polyploid evolution. Molecular Phylogenetics and Evolution 33:197–213.

Evans, B. J., J. C. Morales, M. D. Picker, D. B. Kelley, and D. J. Melnick. 1997. Comparative molecular phylogeography of two Xenopus species, X. gilli and X. laevis, in the southwestern Cape Province, South Africa. Molecular Ecology 6:333–343.

Ferreri, M., W. Qu, and B. Han. 2011. Phylogenetic networks: A tool to display character conflict and demographic history. African Journal of Biotechnology 10:12799–12803.

Frost, D. R. 2021. Amphibian Species of the World: an Online Reference. Version 6.1 (15 November 2021). American Museum of Natural History, New York, USA.

Furman, B. L., A. J. Bewick, T. L. Harrison, E. Greenbaum, V. Gvozdik, C. Kusamba, and B. J. Evans. 2015. Pan-African phylogeography of a model organism, the African clawed frog ‘Xenopus laevis’. Mol Ecol 24:909–925.

Furman, B. L. S., U. J. Dang, B. J. Evans, and G. B. Golding. 2018. Divergent subgenome evolution after allopolyploidization in African clawed frogs (Xenopus). J Evol Biol 31:1945–1958.

Gaeta, R. T. and J. C. Pires. 2010. Homoeologous recombination in allopolyploids: the polyploid ratchet. New Phytologist 186:18–28.

Gaeta, R. T., J. C. Pires, F. Iniguez-Luy, E. Leon, and T. C. Osborn. 2007. Genomic changes in resynthesized Brassica napus and their effect on gene expression and phenotype. Plant Cell 19:3403–3417.

Gelman, A. and J. Hill. 2006. Data Analysis Using Regression and Multilevel/Hierarchical Models (Analytical Methods for Social Research). Cambridge University Press, Cambridge, UK.

Ginal, P., M. Mokhatla, N. Kruger, J. Secondi, A. Herrel, J. Measey, and D. Roedder. 2021. Ecophysiological models for global invaders: Is Europe a big playground for the African clawed frog? Journal of Experimental Zoology 335(1): 158–172.

Green, R. E., et, and al. 2010. A draft sequence of the neandertal genome. Science 328:710–722.

Grohovaz, G. S., B. Harley, and B. Fabran. 1996. Significant mitochondrial DNA sequence divergence in natural populations of Xenopus laevis (Pipidae) from South Africa. Herpetologica 52:247–253.

Gurdon, J. B. 1996. Introductory comments: Xenopus as a laboratory animal. Pp. 3–6 in R. C. Tinsley, and H. R. Kobel, eds. The Biology of Xenopus. Oxford University Press, Oxford.

Gurdon, J. B. and N. Hopwood. 2000. The introduction of Xenopus laevis into developmental biology: of empire, pregnancy testing and ribosomal genes. International Journal of Developmental Biology 44:43–50.

Harland, R. M. and R. M. Grainger. 2011. Xenopus research: Metamorphosed by genetics and genomics. Trends in Genetics 27:507–515.

Ihlow, F., J. Courant, J. Secondi, A. Herrel, R. Rebelo, G. J. Measey, F. Lillo, F. A. De Villiers, S. Vogt, C. De Busschere, T. Backeljau, and D. Rodder. 2016. Impacts of Climate Change on the Global Invasion Potential of the African Clawed Frog Xenopus laevis. PLoS One 11:Article No.: e0154869.

Joshi, N. 2011. Sabre - A barcode dsemultiplexing and trimming tool for FastQ files. GitHub, San Francisco, CA.

Jukes, T. H. and C. R. Cantor. 1969. Evolution of protein molecules. Pp. 21–132 in H. N. Munro, ed. Mammalian Protein Metabolism. Academic Press, New York.

Kobel, H. R. and L. Du Pasquier. 1986. Genetics of polyploid Xenopus. Trends in Genetics 2:310–315.

Kobel, H. R., C. Loumont, and R. C. Tinsley. 1996. The extant species. Pp. 9–33 in R. C. Tinsley, and H. R. Kobel, eds. The Biology of Xenopus. Clarendon Press, Oxford.

Kruger, N., J. Secondi, L. du Preez, A. Herrel, and J. Measey. 2022. Phenotypic variation in Xenopus laevis tadpoles from contrasting climatic regimes is the result of adaptation and plasticity. Oecologica.

Lee, J. and K. Adams. 2020. Global insights into duplicated gene expression and alternative splicing in polyploidBrassica napusunder heat, cold, and drought stress. Plant Genome 13.

Leigh, J. and D. Bryant. 2015. POPART: full-feature software for haplotype network construction. Methods in Ecology and Evolution 6:1110–1116.

Li, H. and R. Durbin. 2009. Fast and accurate short read alignment with Burrows-Wheeler transform. Bioinformatics 25:1754–1760.

Li, H., B. Handsaker, A. Wysoker, T. Fennell, J. Ruan, N. Homer, G. Marth, G. Abecasis, and R. Durbin. 2009. The sequence alignment/map (SAM) format and SAMtools. Bioinformatics 25:2078–2079.

Lillo, F., C. Dufresnes, F. P. Faraone, M. Lo Valvo, and M. Stöck. 2013. Identification and potential origin of invasive clawed frogs Xenopus (Anura: Pipidae) in Sicily based on mitochondrial and nuclear DNA. Italian Journal of Zoology 80:566–573.

Liu, S., Y. Liu, X. Yang, C. Tong, D. Edwards, I. A. P. Parkin, M. Zhao, J. Ma, J. Yu, S. Huang, X. Wang, J. Wang, K. Lu, Z. Fang, I. Bancroft, T.-J. Yang, Q. Hu, X. Wang, Z. Yue, H. Li, L. Yang, J. Wu, Q. Zhou, W. Wang, G. J. King, J. C. Pires, C. Lu, Z. Wu, P. Sampath, Z. Wang, H. Guo, S. Pan, L. Yang, J. Min, D. Zhang, D. Jin, W. Li, H. Belcram, J. Tu, M. Guan, C. Qi, D. Du, J. Li, L. Jiang, J. Batley, A. G. Sharpe, B.-S. Park, P. Ruperao, F. Cheng, N. E. Waminal, Y. Huang, C. Dong, L. Wang, J. Li, Z. Hu, M. Zhuang, Y. Huang, J. Huang, J. Shi, D. Mei, J. Liu, T.-H. Lee, J. Wang, H. Jin, Z. Li, X. Li, J. Zhang, L. Xiao, Y. Zhou, Z. Liu, X. Liu, R. Qin, X. Tang, W. Liu, Y. Wang, Y. Zhang, J. Lee, H. H. Kim, F. Denoeud, X. Xu, X. Liang, W. Hua, X. Wang, J. Wang, B. Chalhoub, and A. H. Paterson. 2014. The Brassica oleracea genome reveals the asymmetrical evolution of polyploid genomes. Nature Communications 5:Article No.: 3930.

Liu, Y., J. Wang, W. Ge, Z. Wang, Y. Li, N. Yang, S. Sun, L. Zhang, and X. Wang. 2017. Two Highly Similar Poplar Paleo-subgenomes Suggest an Autotetraploid Ancestor of Salicaceae Plants. Frontiers in Plant Science 8:Article No.: 571.

Louppe, V., B. Leroy, A. Herrel, and G. Veron. 2020. The globally invasive small Indian mongoose Urva auropunctata is likely to spread with climate change. Scientific Reports 10:Article No.: 7461.

Lovell, J. T., A. H. MacQueen, S. Mamidi, J. Bonnette, J. Jenkins, J. D. Napier, A. Sreedasyam, A. Healey, A. Session, S. Shu, K. Barry, S. Bonos, L. Boston, C. Daum, S. Deshpande, A. Ewing, P. P. Grabowski, T. Haque, M. Harrison, J. Jiang, D. Kudrna, A. Lipzen, T. H. Pendergast, C. Plott, P. Qi, C. A. Saski, E. Shakirov, V D. Sims, M. Sharma, R. Sharma, A. Stewart, V. R. Singan, Y. Tang, S. Thibivillier, J. Webber, X. Weng, M. Williams, G. A. Wu, Y. Yoshinaga, M. Zane, L. Zhang, J. Zhang, K. D. Behrman, A. R. Boe, P. A. Fay, F. B. Fritschi, J. D. Jastrow, J. Lloyd-Reilley, J. M. Martinez-Reyna, R. Matamala, R. B. Mitchell, F. M. Rouquette, P. Ronald, M. Saha, C. M. Tobias, M. Udvardi, R. A. Wing, Y. Wu, L. E. Bartley, M. Casler, K. M. Devos, D. B. Lowry, D. S. Rokhsar, J. Grimwood, T. E. Juenger, and J. Schmutz. 2021. Genomic mechanisms of climate adaptation in polyploid bioenergy switchgrass. Nature (London) 590:438–444.

Martin, M. 2011. Cutadapt removes adapter sequences from high-throughput sequencing reads. EMBnet. journal 17:pp–10.

Martin, S. 2021. Genomics_general repository. GitHub, San Francisco, CA.

Matsuda, Y., Y. Uno, M. Kondo, M. J. Gilchrist, A. M. Zorn, D. S. Rokhsar, M. Schmid, and M. Taira. 2015. A new nomenclature of Xenopus laevis chromosomes based on the phylogenetic relationship to Silurana/Xenopus tropicalis. Cytogenetic and Genome Research 145:187–191.

McKenna, A., M. Hanna, E. Banks, A. Sivachenko, K. Cibulskis, A. Kernytsky, K. Garimella, D. Altshuler, S. Gabriel, M. Daly, and M. A. DePristo. 2010. The Genome Analysis Toolkit: a MapReduce framework for analyzing next-generation DNA sequencing data. Genome Research 20:1297–1303.

Measey, G. J. 2004. Species account: Xenopus laevis (Daudin 1802). Pp. 266-267 in L. R. Minter, M. Burger, J. A. Harrison, and H. H. Braack, eds. Atlas and Red Data Book of the Frogs of South Africa, Lesotho and Swaziland. Smithsonian Institution Press, Washington D. C., USA.

Measey, G. J. and A. Channing. 2003. Phylogeography of the genus Xenopus in southern Africa. Amphibia-Reptilia 24:321–330.

Measey, G. J., D. Rodder, S. L. Green, R. Kobayashi, F. Lillo, R. Rebelo, and J.-M. Thirion. 2012. Ongoing invasions of the African clawed frog, Xenopus laevis: a global review. Biological Invasions 14:2255–2270.

Measey, J. 2016. Overland movement in African clawed frogs (Xenopus laevis): a systematic review. PeerJ 4:Article No.: e2474.

Measey, J. 2017. Where do African clawed frogs come from? An analysis of trade in live Xenopus laevis imported into the USA. Salamandra 53:398–404.

Measey, J., S. J. Davies, G. Vimercati, A. Rebelo, W. Schmidt, and A. Turner. 2017. Invasive amphibians in southern Africa: A review of invasion pathways. Bothalia 47:Article No.: a2117.

Mei, W., S. Liu, J. C. Schnable, C.-T. Yeh, N. M. Springer, P. S. Schnable, and W. B. Barbazuk. 2017. A Comprehensive Analysis of Alternative Splicing in Paleopolyploid Maize. Frontiers in Plant Science 8:Article No.: 694.

Minh, B. Q., M. A. Nguyen, and A. von Haeseler. 2013. Ultrafast approximation for phylogenetic bootstrap. Molecular Biology and Evolution 30:1188–1195.

Mucina, L. and M. C. Rutherford. 2006. The vegetation of South Africa, Lesotho and Swaziland, Strelitzia 19. South African National Biodiversity Institute, Pretoria, South Africa.

Nguyen, L. T., H. A. Schmidt, A. von Haeseler, and B. Q. Minh. 2015. IQ-TREE: A fast and effective stochastic algorithm for estimating maximum-likelihood phylogenies. Molecular Biology and Evolution 32:268–274.

Ortiz, E. M. 2019. vcf2phylip v2.0: convert a VCF matrix into several matrix formats for phylogenetic analysis.

Otto, S. P. and J. Whitton. 2000. Polyploid incidence and evolution. Annual Review of Genetics 34:401–437.

Parisod, C., C. Definod, A. Sarr, N. Arrigo, and F. Felber. 2013. Genome-specific introgression between wheat and its wild relative Aegilops triuncialis. Journal of Evolutionary Biology 26:223–228.

Patterson, N., P. Moorjani, Y. Luo, S. Mallick, N. Rohland, Y. Zhan, T. Genschoreck, T. Webster, and D. Reich. 2012. Ancient admixture in human history. Genetics 192:1065–1093.

Peralta-Garcia, A., J. H. Valdez-Villavicencio, and P. Galina-Tessaro. 2014. African clawed frog (Xenopus laevis) in Baja California: a confirmed population and possible ongoing invasion in mexican watersheds. Southwestern Naturalist 59:431–434.

Rebelo, R., P. Amaral, M. Bernardes, J. Oliveira, P. Pinheiro, and D. Leitao. 2010. Xenopus laevis (Daudin, 1802), a new exotic amphibian in Portugal. Biological Invasions 12:3383–3387.

Schiavinato, M., A. Bodrug-Schepers, J. C. Dohm, and H. Himmelbauer. 2021. Subgenome evolution in allotetraploid plants. Plant Journal 106.

Session, A. M., Y. Uno, T. Kwon, J. A. Chapman, A. Toyoda, S. Takahashi, A. Fukui, A. Hikosaka, A. Suzuki, M. Kondo, S. J. van Heeringen, I. Quigley, S. Heinz, H. Ogino, H. Ochi, U. Hellsten, J. B. Lyons, O. Simakov, N. Putnam, J. Stites, Y. Kuroki, T. Tanaka, T. Michiue, M. Watanabe, O. Bogdanovic, R. Lister, G. Georgiou, S. S. Paranjpe, I. van Kruijsbergen, S. Shu, J. Carlson, T. Kinoshita, Y. Ohta, S. Mawaribuchi, J. Jenkins, J. Grimwood, J. Schmutz, T. Mitros, S. V. Mozaffari, Y. Suzuki, Y. Haramoto, T. S. Yamamoto, C. Takagi, R. Heald, K. Miller, C. Haudenschild, J. Kitzman, T. Nakayama, Y. Izutsu, J. Robert, J. Fortriede, K. Burns, V. Lotay, K. Karimi, Y. Yasuoka, D. S. Dichmann, M. F. Flajnik, D. W. Houston, J. Shendure, L. DuPasquier, P. D. Vize, A. M. Zorn, M. Ito, E. M. Marcotte, J. B. Wallingford, Y. Ito, M. Asashima, N. Ueno, Y. Matsuda, G. J. Veenstra, A. Fujiyama, R. M. Harland, M. Taira, and D. S. Rokhsar. 2016. Genome evolution in the allotetraploid frog Xenopus laevis. Nature 538:336–343.

Sharbrough, J., J. L. Conover, M. F. Gyorfy, C. E. Grover, E. R. Miller, J. F. Wendel, and D. B. Sloan. 2021. Global patterns of subgenome evolution in organelle-targeted genes of six allotetraploid angiosperms. BioRxiv in review.

Soares, N. R., M. Mollinari, G. K. Oliveira, G. S. Pereira, and M. L. C. Vieira. 2021. Meiosis in Polyploids and Implications for Genetic Mapping: A Review. Genes 12:Article No.: 1517.

Swinscow, T. D. V. and M. J. Campbell. 2002. Statistics at Square One. BMJ, London.

Swofford, D. L. 2002. Phylogenetic analysis using parsimony (* and other methods). Version 4. Sinauer Associates, Sunderland.

Tinsley, R. C., C. Loumont, and H. R. Kobel. 1996. Geographical distribution and ecology. Pp. 35–59 in R. C. Tinsley, and H. R. Kobel, eds. The Biology of Xenopus. Clarendon Press, Oxford.

Tolley, K.A., Bowie, R.C.K., Measey, G.J., Price, B.W. and Forest, F. 2014. The shifting landscape of genes since the Pliocene: terrestrial phylogeography in the Greater Cape Floristic Region In: Fynbos: ecology, evolution, and conservation of a megadiverse region. (eds. Allsopp, N., Colville, J.F., Verboom, G.A.) pp 143–163. Oxford University Press.

Van de Peer, Y., S. Maere, and A. Meyer. 2009. The evolutionary significance of ancient genome duplications. Nature Reviews Genetics 10:725–732.

van Sittert, L. and J. Measey. 2016. Historical perspectives on global exports and research of African clawed frogs (Xenopus laevis). Transactions of the Royal Society of South Africa 71:157–166.

Vimercati, G., N. Kruger, and J. Secondi. 2021. Land cover, individual’s age and spatial sorting shape landscape resistance in the invasive frog Xenopus laevis. Journal of Animal Ecology 90.

Wagener, C., N. Kruger, and J. Measey. 2021. Progeny of Xenopus laevis from attitudinal extremes display adaptive physiological performance. Journal of Experimental Biology 224.

Wang, S., Y. Hong, and J. Measey. 2019. An established population of African clawed frogs, Xenopus laevis (Daudin, 1802), in mainland China. BioInvasions Records 8:457–464.

Wei, Z., M. Wang, S. Chang, C. Wu, P. Liu, J. Meng, and J. Zou. 2016. Introgressing subgenome components from Brassica rapa and B. carinata to B. juncea for broadening Its Genetic Base and Exploring Intersubgenomic Heterosis. Frontiers in Plant Science 7:Article No.: 1677.

Weisman, A. I. and C. W. Coates. 1941. The frog test (Xenopus laevis) as a rapid diagnostic test for early pregnancy. Endocrinology 28:141–142.

Weldon, C., A. De Villiers, and L. Du Preez. 2007. Quantification of the trade in Xenopus laevis from South Africa, with implications for biodiversity conservation. African Journal of Herpetology 56:77–83.

Wolfe, K. H. 2001. Yesterdays’s polyploids and the mystery of diploidization. Nature Reviews Genetics 2:333–341.

Wu, H., Q. Yu, J.-H. Ran, and X.-Q. Wang. 2021. Unbiased Subgenome Evolution in Allotetraploid Species of Ephedra and Its Implications for the Evolution of Large Genomes in Gymnosperms. Genome Biology and Evolution 13.

Yu, K., M. Feng, G. Yang, L. Sun, Z. Qin, J. Cao, J. Wen, H. Li, Y. Zhou, X. Chen, H. Peng, Y. Yao, Z. Hu, W. Guo, Q. Sun, Z. Ni, K. Adams, and M. Xin. 2020. Changes in Alternative Splicing in Response to Domestication and Polyploidization in Wheat. Plant Physiology 184:1955–1968.

Zhao, Y., L. Dong, C. Jiang, X. Wang, J. Xie, M. A. R. Rashid, Y. Liu, M. Li, Z. Bu, H. Wang, X. Ma, S. Sun, X. Wang, C. Bo, T. Zhou, and L. Kong. 2020. Distinct nucleotide patterns among three subgenomes of bread wheat and their potential origins during domestication after allopolyploidization. BMC Biology 18: 188.

Zheng, X., D. Levine, J. Shen, S. M. Gogarten, C. Laurie, and B. S. Weir. 2012. A high-performance computing toolset for relatedness and principal component analysis of SNP data. Bioinformatics 28:3326–3328.

